# Predicting Morphological Disparities in Sea Urchin Skeleton Growth and Form

**DOI:** 10.1101/133900

**Authors:** Maria Abou Chakra, Miroslav Lovric, Jonathon Stone

**Affiliations:** Max Planck Institute for Evolutionary Biology, Department of Evolutionary Theory, Plön, SH 24306, Germany.; McMaster University, Department of Biology, 1280 Main Street West, Hamilton, Ontario L8S 4K1, Canada; McMaster University, Mathematics & Statistics Department, 1280 Main Street West, Hamilton, Ontario L8S 4M1, Canada; McMaster University, Origins Institute, 1280 Main Street West, Hamilton, Ontario L8S 4M1, Canada; McMaster University, SHARCNet, 1280 Main Street West, Hamilton, Ontario L8S 4M1, Canada

**Keywords:** Theoretical Morphology, Development, Sea Urchins, Paleontology, Skeleton, Morphological Disparity

## Abstract

Sea urchins exhibit among their many species remarkable diversity in skeleton form (e.g., from spheroid to discoid shapes). However, we still do not understand how some related species show distinct morphologies despite inherent similarities at the genetic level. For this, we use theoretical morphology to disentangle the ontogenic processes that play a role in skeletal growth and form. We developed a model that simulates these processes involved and predicted trajectory obtaining 94% and 77% accuracies. We then use the model to understand how morphologies evolved by exploring the individual effects of three structures (ambulacral column, plate number, and polar regions). These structures have changed over evolutionary time and trends indicate they may influence skeleton shape, specifically height–to-diameter ratio, h:d. Our simulations confirm the trend observed but also show how changes in the attributes affect shape; we show that widening the ambulacral column or increasing plate number in columns produces a decrease in h:d (flattening); whereas increasing apical system radius to column length ratio produces an increase in h:d (gloublar shape). Computer simulated h:d matched h:d measured from real specimens. Our findings provide the first explanation of how small changes in these structures can create the diversity in skeletal morphologies.

## Introduction

Researchers still puzzle over how some related species show distinct morphologies despite inherent similarities at the genetic level, the genotypic-phenotypic paradox. Humans and chimpanzee species, for instance, exhibit distinct morphologies despite the similarity at gene and protein levels^1^. This puzzle extends to larger taxonomic groups, such as the echinoderm class Echinoidea, where extremely different skeletal morphologies are manifested from the same structural attributes.

Echinoid skeletons (tests) exhibit pentameric symmetry and the vast morphological disparity among species can be observed throughout the fossil record, which dates to the Late Ordovician^2,3^. The earliest sea urchins (Paleozoic echinoids) exhibited spheroid body forms; however, by the Jurassic, after members in the Irregularia (sand dollars, sea biscuits, and heart urchins) first appeared, discoid-shaped, bottle-shaped, and even heart-shaped body forms had evolved^2-6^. The diversity has elicited equally diverse developmental, evolutionary, and adaptationist explanations among morphologists^3-5,7-14^. Explaining the disparity, however, specifically how it is effected during development and growth, remains challenging. One reason for the persistent challenge resides in disentangling the 5 underlying ontogenic processes, plate growth, plate addition, plate interaction, plate gapping, and visceral growth^15^, which are interrelated and occur simultaneously.

All echinoid (sea urchin) tests comprise plates. The plates are produced and translated within growth zones, the five-fold repeating unit constituting the pentaradial test^10,12,16^. Each growth zone extends from the aboral surface (containing the apical system) to the oral surface (containing the peristome). Plates occupy three regions (Fig. 1): the apical system (comprising genital plates and ocular plates), corona (comprising ambulacral plates and interambulacral plates), and peristome (comprising buccal plates). During growth, new plates are added at the apical system and old plates interlock and change size; plates must separate from one another to increase or decrease in size^11,17-19^.

**Figure 1.**
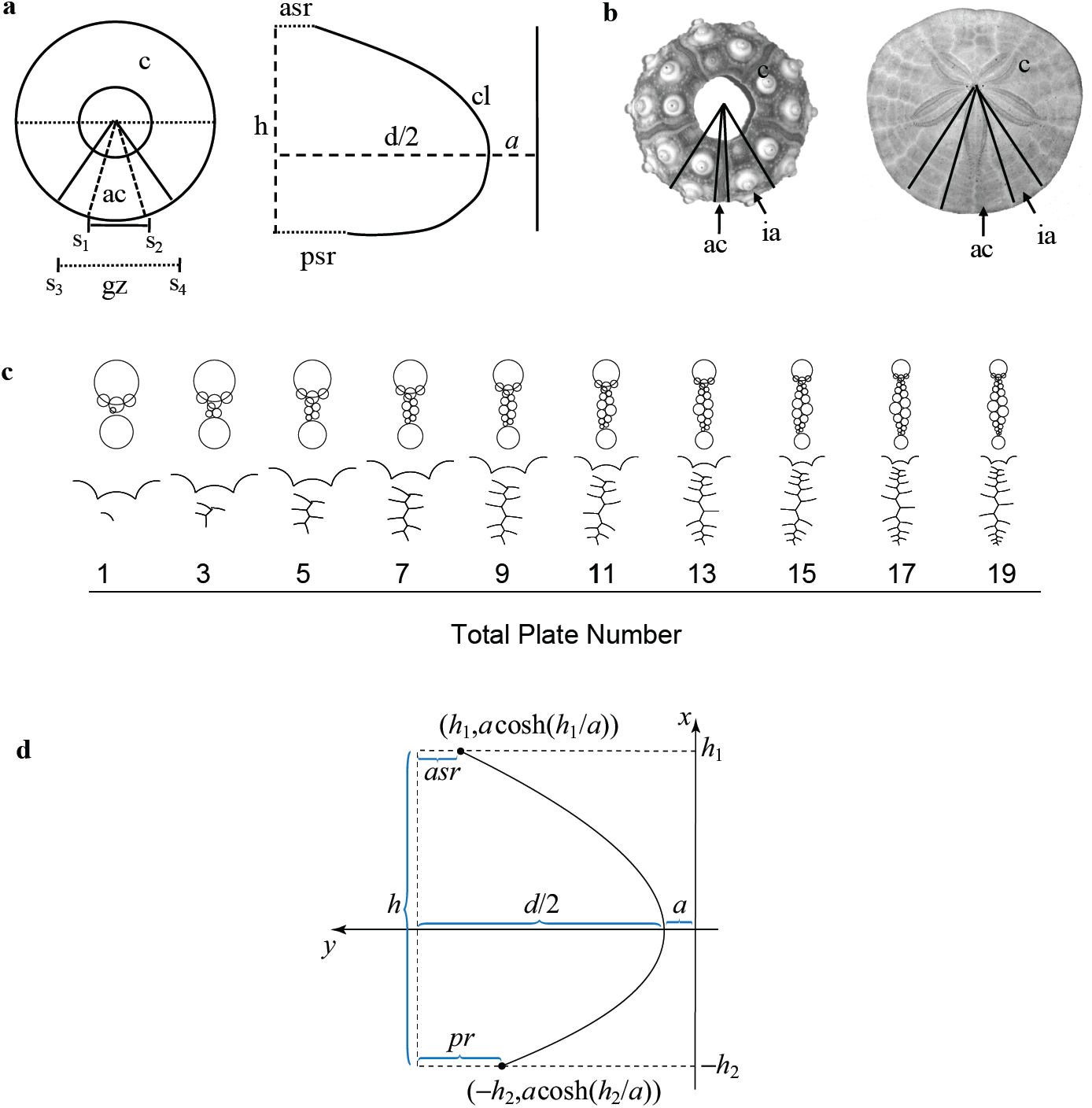
Schematic representation for an echinoid test. a) structural attributes: corona, c, ambulacral column, ac (measured as the angle spanning line segment s_1_-s_2_), growth zone, gz (measured as the angle spanning line segment s_3_-s_4_), apical system radius, asr, column length, cl, ambitus diameter, d, peristome radius, pr, interambulacral column, ia, and test height, h. b) aboral surface for *Eucidaris thouarsii* (left) and *Dendraster excentricus* (right), displaying the size difference between the columns (am and ia) within a growth zone. c) Simulating the growth of an ambulacral column shown over time as new plates are added. Parameters used ac=40°, as=0.015mm, tpn=20, p:as=0.75, and asr:cl=0.25 d) Relationship between catenary curve and echinoid test, apical system radius and peristome radius, height (*= h*_*1*_ *+ h*_*2*_), ambitus diameter to predict *a* using *a* + d/2 – asr = *a* cosh (h_1_ / *a*) and *a* + d / 2 – p = *a* cosh (h_2_ / *a*).

Skeletons exhibit patterns and shapes that can be captured using elementary principles and mathematical techniques^20,21^. Over the past century, 9 published theoretical models^9,15,20,22-28^ have been developed to explain echinoid test growth^29^. Holotestoid^15161717^ is a theoretical computational model that incorporates the mathematical and physical principles involved in coalescing bubbles, close-packing circles, and, as developed herein, catenaries to emulate each process involved in test growth and form. Coalescing bubbles, close-packing circles, and catenaries are well established as solutions to minimal surface problems^15,30-34^ and are associated with natural patterns^20,35^. We herein use Holotestoid to develop in silico growth zones. We use the model to show how discoid, adult body forms can be grown from a spheroid, juvenile body forms. We predict height to diameter ratios, h:d and compare computer-simulated species-specific h:d to measurements from real specimens.

## Materials and Methods

### Empirical Measurements

Specimens of *Eucidaris thouarsii* (n=6) were obtained from the California Academy of Sciences collection, San Francisco, CA, USA; specimens of *Arbacia punctulata* (n=33) were obtained from Gulf Specimen Marine Laboratory, Panacea, FL, USA; specimens of *Lytechinus variegatus* (n=70) and *Mellita quinquiesperforata* (n=10) were obtained from the Marine Biological Laboratory, Woods Hole, MA, USA; and specimens of *Dendraster excentricus* (n=49) and *Strongylocentrotus franciscanus* (n=14) were obtained from Westwind Sealab Supplies, Victoria, BC, Canada.

We chose these 6 morphologically disparate species from the Cidaroida (i.e., *Eucidaris thouarsii*), Stirodonta (i.e., *Arbacia punctulata*), Camarodonta (i.e., *Lytechinus variegatus* and *Strongylocentrotus franciscanus*), and Irregularia (i.e., *Dendraster excentricus* and *Mellita quinquiesperforata*), thus providing a variety of taxonomic samples. Measurements were performed using a Vernier calliper and flexible measuring tape on despined, eviscerated, and cleaned tests. For each specimen, we measured ambulacral column angle, ac, apical system, as, column length, cl, diameter, d, height, h, and peristome p (Fig. 1a; Table 1).

**Table 1.** *E. thouarsii* (Et)*, A. punctulata* (Ap)*, L. variegatus* (Lv)*, S. franciscanus* (Sf)*, M. quinquiesperforata* (Mq), and *D. excentricus* (De) measurements; ambulacral column angle (ac), apical system radius (apr), column length (cl), test diameter (d), test height (h), and peristome radius (pr). Measurements are presented as ac, asr:cl, pr:asr, d:cl, and h:d ± standard error.

|  | ac angle | asr:cl | pr:asr | d:cl | h:d |
| --- | --- | --- | --- | --- | --- |
| Et | 15° ± 1 | 0.18 ± 0.02 | 1.33 ± 0.24 | 1.00 ± 0.07 | 0.57 ± 0.05 |
| Ap | 20° ± 2 | 0.14 ± 0.01 | 1.97 ± 0.21 | 1.10 ± 0.05 | 0.53 ± 0.03 |
| Lv | 27° ± 2 | 0.08 ± 0.02 | 1.77 ± 0.17 | 0.98 ± 0.22 | 0.60 ± 0.05 |
| Sf | 31° ± 3 | 0.09 ± 0.01 | 1.86 ± 0.16 | 1.02 ± 0.08 | 0.44 ± 0.05 |
| Mq | 39° ± 3 | 0.04 ± 0.01 | 0.53 ± 0.06 | 1.07 ± 0.04 | 0.10 ± 0.01 |
| De | 41° ± 3 | 0.03 ± 0.01 | 0.84 ± 0.10 | 1.03 ± 0.16 | 0.15 ± 0.02 |

## Model

The computational model Holotestoid^15^ was designed on principles that capture growth in regular (i.e., spheroid) sea urchins. The principles emulate computationally the 5 ontogenic processes (the original code is available in a data repository^15^).

The first process, plate growth, increases or decreases plate size, with plate size determined on the basis of their location in the growth zone and relative distance from a polar region (apical system, as, or peristome, p; Fig. 1b).

The second process, plate addition, inserts new plates apically at a nucleation point situated next to an ocular plate. Plates are added sequentially, alternating between the left and right side adjacent to each ocular plate.

The third process, plate interaction, involves an analogy in which individual plates are likened to bubbles (circles in two-dimensions) to predict the interfaces and shapes adopted between plates in a column^15,20,22,27,36^.

The fourth process, plate gapping, separates plates (circles) in a close-packing configuration (with no overlaps and minimal gaps), emulating collagen fibre loosening to create gaps for new plate addition and peripheral calcite deposition to occur^17^. The computational model arranges unequal-sized circles in a triangular circle-close-packing configuration to mimic natural gaping.

These four processes result in a growth model where plates are added over time and column length develops according to ambulacra column width (Fig. 1c). The simulations produce graphically growth zones in two dimensions. However, since h:d traditionally is used to describe body forms^4,5,7-9,13,14,37^, we now introduce to Holotestoid the fifth process, visceral growth. This process describes the integrated effects imparted by somatic growth onto test structures^9,37,38^. In some previous models, visceral growth was associated with mathematical curves that describe test outline shapes. The Young-Laplace equation was considered previously by Thompson^20^, in the liquid drop model, and Ellers^26^, in the membrane model (the membrane model was successful for some regular echinoids but was limited in its application, simulating inaccurately outline shapes for tests characterising Cidaroida and Irregularia). We designed the visceral growth module in the computational model to be flexible enough to implement any mathematical function that represents reasonably visceral growth (e.g., parabolic or hyperbolic); however, we selected curves (catenaries, or catenary curves) that utilise data extracted easily by measurement, such as asr, cl, pr, and d, and are a (closely approximate) consequence of the dynamics of the test growth.

Catenaries describe the shape that is assumed by an inextensible but flexible chain that hangs freely from two fixed points (this shape can model other objects, such as dental arches^26,43^). A catenary curve models the balance between two forces in echinoid tests (Fig. S1): the tension between plates and the horizontal force acting in the outward direction. We analogised each plate as a separate link in a chain with the tension force pulling plates apart^26,38^ – this analogy is based on the idea that plates can contract and relax, as some are sutured together^8,13,26^.

A catenary curve is represented as a function with the general form:

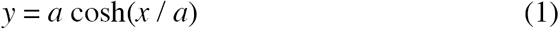

with (0, *a*) as its lowest point (Fig. 1d). In the model, x represents height (*h*) and is given by *h = h*_*1*_ *+ h*_*2*_. When *x = h*_*1*_, then *y* = *a + d /* 2 *-asr* and thus equation (1) implies that

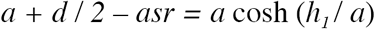

Likewise, when *x* = -*h*_*2*_, then *y* = *a + d / 2 - pr*, and equation (1) implies that

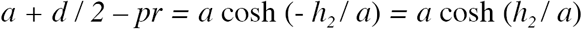

We solve this system of three equations for *a*, *h*_*1*_ and *h*_*2*,_ using numeric methods. Calculating the value for *a* needed in equation (1) requires knowing test height, h, ambitus diameter, d, apical system radius, asr, and peristome system radius, pr (Fig 1). As described previously, the first four processes involve values for asr, pr, and cl. We lack a predictor for diameter, d; thus, we need to use *cl* to estimate test height,

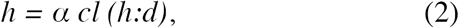

where h:d is taken as a parameter and *d = α cl*, where *α* is a constant. In the model, we combined equations (1) and (2) to predict test shape by determining the constant *a* in equation (1) (Fig. 1d). We created a function that cycles through a range of possible *h:d* ratios (from 0.1 to 0.9, in 0.0001-step increments), predicting curvature for each incremental value. As is the case with applying any function, suitable justification is required; we therefore validated model predictions with measurements taken from real specimens.

Theoretically, two extreme morphologies may be realized as column length increases in the limit to infinity, extremely flattened tests or extremely domed tests. Based on the 6 species measured (Table 1), typically h:d < 1. Although h:d > 1 theoretically is possible, our empirical data revealed that the average d:cl is 1.0 ± 0.1 for all species (Table 1, Fig S2). As column length increases, diameter increases under a fixed d:cl constraint, thus restricting tests to h:d<1. This assumption can be relaxed in the future, but for now it provides a straightforward transition between 2-dimensional and 3-dimensional perspectives.

The visceral module requires 3 parameters: column length, *cl*, apical system radius, *asr*, and peristome radius, *pr,* to predict test height, h. Values for these can be extracted from the model or measured from real specimens. We used all the measured attributes (Table 1) as parameters and compared measured d and h with values predicted from the module, obtaining 94% and 77% accuracies (Fig. S2). This confirms that catenaries can be used to predict accurately h:d for regular echinoids. We then combined the visceral growth module with the computational model, Holotestoid.

### Simulations

Combining the 5 processes, we simulated sea urchin growth zones with 6 parameters: growth zone angle (gz, Fig. 1a), ambulacral column angle (ac, Fig. 1a), total ambulacral plate number (tpn), apical system radius (asr, Fig. 1b), peristome-to-apical system ratio (p:as), and apical system-to-column length ratio (asr:cl). The p:as and asr:cl are maintained throughout each simulation unless asr:cl=0, which entails that polar region sizes remain fixed while columns grew.

Parameter value ranges for all simulations were chosen to encompass measured data for real specimens (Table 1). We assumed that growth zones were equivalent in arc length, so growth zone angle was fixed, gz = 72°, and initial apical system diameter was set arbitrarily at 1 mm.

## Results and Discussion

Simulations to predict species specific h:d were conducted using parameter ranges encompassing measured data from real specimens (Table1). We measured ambitus diameter, d, test height, h, and column length, cl, from 6 disparate species (Table 1): 4 regular (i.e., *Eucidaris thouarsii, Arbacia punctulata*, *Lytechinus variegatus* and *Strongylocentrotus franciscanus*) and 2 irregular (i.e., *Dendraster excentricus* and *Mellita quinquiesperforata*). The simulated developmental trajectories for each species spanned from the juvenile stage to an adult stage. For each new plate addition and thus skeleton growth increment, the model predicted a h:d value. For each simulated representative for a particular species, reasonable h:d (Fig. 2), within the measured range, were predicted: the *E. thouarsii* repersentative h:d reached 0.63±0.03, which falls within measured h:d range 0.57 ± 0.05 (Fig. 2a). The *A. punctulata* representative h:d reached 0.56±0.04, which falls within measured h:d range 0.53 ± 0.03 (Fig. 2b). The *L. variegatus* representative h:d reached 0.59±0.07, which falls within measured h:d range 0.60 ± 0.05 (Fig. 3c). The *S. franciscanus* representative h:d reached 0.45±0.05, which falls within measured h:d range 0.44 ± 0.05 (Fig. 3d). The *M. quinquiesperforata* representative h:d reached 0.14±0.02, close to the measured h:d range 0.10 ± 0.01 (Fig. 3e). And the *D. excentricus* representative h:d reached 0.15 ± 0.02, which falls within measured h:d range 0.12±0.01 (Fig. 3f).

**Figure 2.**
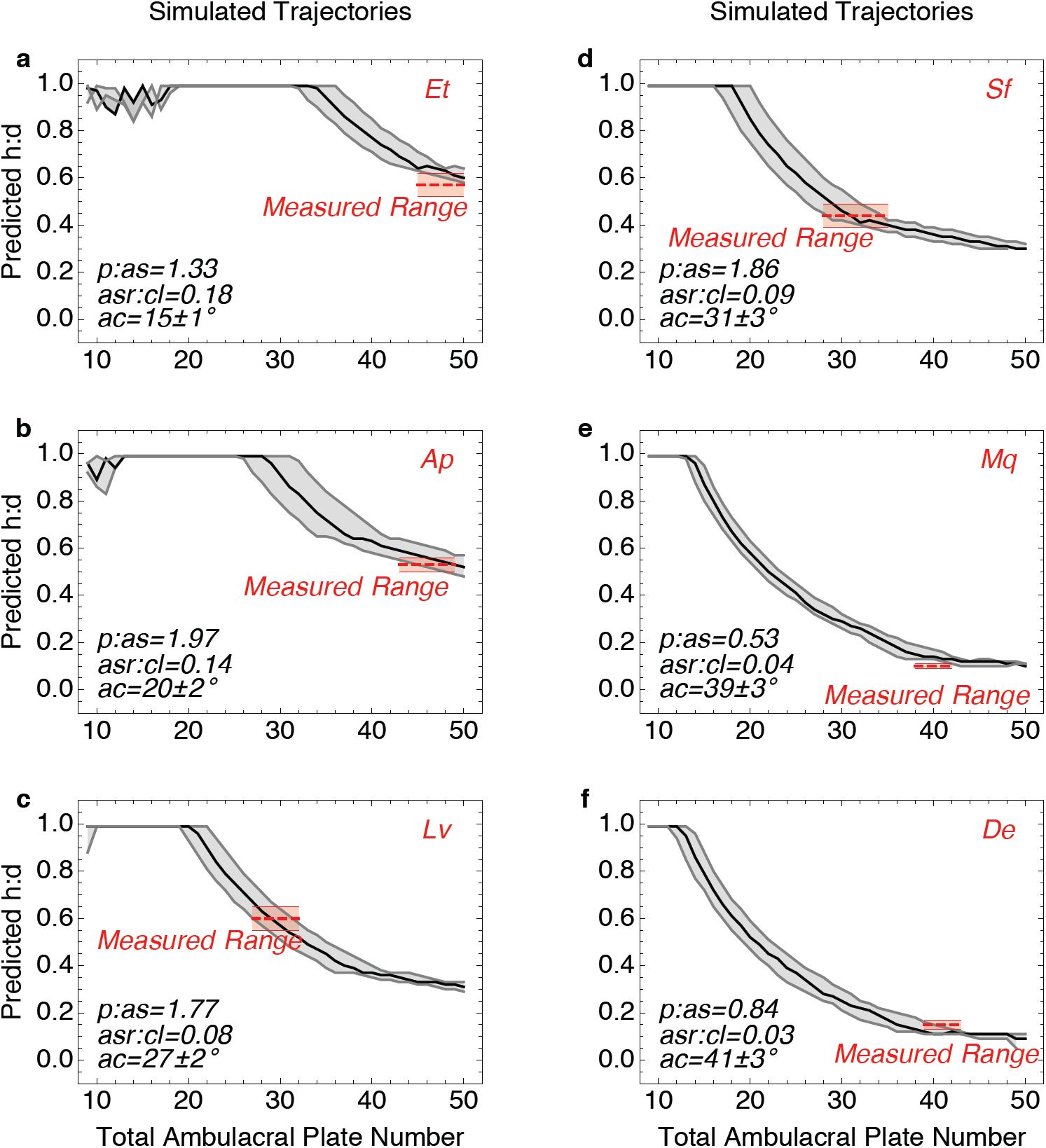
Simulations of growth trajectories for using species specific parameters. We impose measured range for height to diameter ratio, h:d (dashed red line), from real specimens onto and the predicted h:d (gray curves and black curves). We used the parameter values measured from representative specimens from six species (Table 1): a) *E. thouarsii* (Et), b) *A. punctulata* (Ap), c) *L. variegatus* (Lv), d) *S. franciscanus* (Sf), e) *M. quinquiesperforata* (Mq), f) *D. excentricus* (De). General parameters used ap=0.015mm, tpn=50, gz=72°,

**Figure 3.**
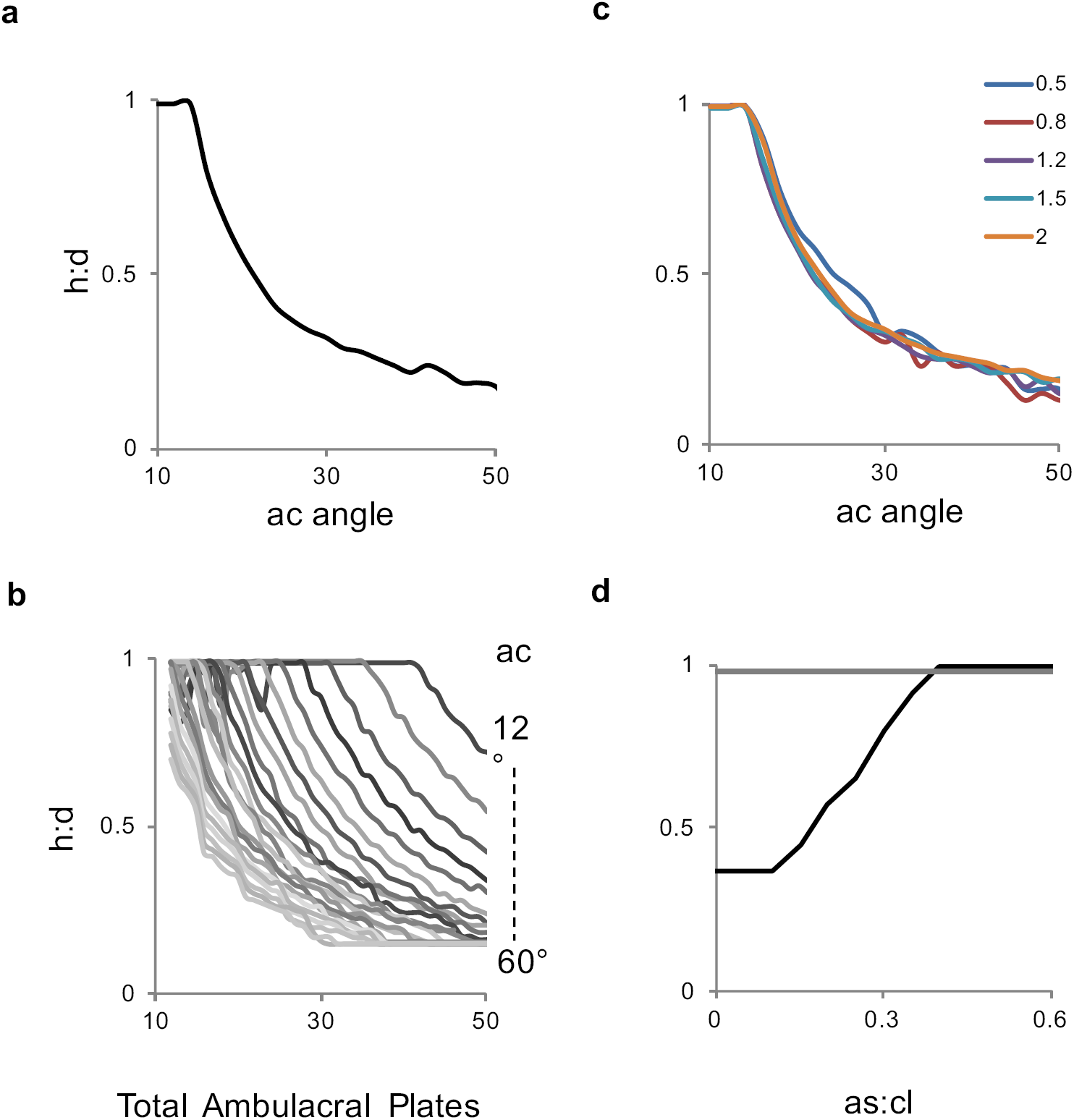
Simulation predicting effects on height to diameter ratio, h:d, with respect to a) am: an increase caused h:d to decrease; b) as total ambulacral plate number, tpn, in a column increased to 50 the h:d decreased, with trends shown as am increased from 12° to 60°. c) The persitome to apical system, p:as, ratio values 0.5, 0.8, 1, 1.2, 1.5, and 2 imparted no effect on h:d (all six curves overlapped and exhibited the same trend as in a); d) An increase in the apical system to column length, ap:cl, ratio caused an increase in h:d, for instance a simulation with a wide ambulacral column, am=40° (black curve), started of flat, low h:d ratio and increase as ap:cl also increase. However tests with a thin ambulacral column, am = 8° already were spheroidal and couldn’t increase any further in h:d (grey curve). Parameters values used unless stated explicitly were asr = 0.05mm, tpn = 25 / 50, p:as = 1, and asr:cl = 0.

We explored effects from the structural attributes ambulacral column width, total plate number, and polar region sizes (apical system radius and peristome radius) on h:d. Simulations reveal that increasing ambulacral column angle, am, from 10° to 64° decreases h:d from 0.99 to 0.1 (Fig. 3a). This shows that increasing ambulacra column width can lead to flattened tests, which is consistent with morphological observations across the post-Palaeozoic record^4^.

Similar effects were observed when total plate number was varied. Simulations revealed that increasing total plate number decreases h:d from 0.99 to 0.16 (Fig. 2b). This entails that changes in plate number can influence overall test shape – increasing tpn can lead to flattened tests. Increasing column angle and plate number can impart compounding effects; for example, the flattening rate changed: for ac=12°, h:d started to decrease at tpn=44, whereas, for ac=60°, h:d started to decrease at tpn=8 (Fig. 3b). We also observed that, for all am, tpn < 10 always produced h:d > 0.9, suggesting a possible explanation for the test shape similarity observed across newly metamorphosed juveniles^17-19,39^, which comprise a small plate number and are characterized by spheroid shapes.

Contrastingly, simulations showed that apical system and peristome sizes, themselves, have no affect on test h:d. Increasing ap from 0.05 to 50 produced no change in h:d, which remained constant at 0.16; increasing ps:ap from 0.5 to 2 similarly produced no net effect on h:d (Fig. 2c). On the basis of these results, we infer that, under particular conditions (*e*.*g*., no growth in the polar regions), h:d is impacted by the structural attributes ambulacral column width and plate number rather than polar region sizes.

Computer simulations revealed that increasing asr:cl from 0 to 0.6 increased h:d from 0.37 to 0.99 (Fig. 2d, black curve). The asr:cl imparted greater influence on columns produced by large am (Fig. 2d, black curve) compared to small am (Fig. 2d, grey curve). From these results, we infer that test h:d is influenced by multiple, even simultaneous, changes among the structural attributes ambulacral column width, plate number, and polar region sizes.

## Conclusion

We present and validate a model that simulates growth zones in sea urchins. We used the model to explore how morphological diversity can be achieved across echinoid groups by changing growth trajectories. We explore how changes to particular structural attributes can produce discoid from spheroid body forms. Our results provide an explanation for how different h:d are achieved in different major echinoid groups and furthermore show how those h:d may be realized throughout development in individuals. Researchers historically have noted that adult h:d is species-specific^4,7,24,40^. This is the first study to show how such h:d may be sustained through balanced growth among the structural attributes comprising tests (ambulacral column width, total plate number, polar region sizes). Our results also suggest a possible explanation for the test shape similarity observed across newly metamorphosed juveniles^17-19,39^.

This study provides an explanation for how the disparity observed between regular echinoids (*e*.*g*., sea urchins) and irregular echinoids (*e*.*g*., sand dollars) evolved. We infer that flattened tests were effected by increases in the ambulacral column width and decreases in apical system radius to column length ratio and peristome radius to column length ratio in irregular echinoids in comparison to regular echinoids. Although this study involved only 6 species, the findings constitute an essential step toward understanding morphological diversity seem across echinoid taxa. The next step would be to expand the analysis to more species. Increasing the predictive accuracy of the model may prove valuable to the unresolved origin of the ancestral shape which is a single fossil specimen ^2,3,27,40-42^.

## Acknowledgments

We thank, F. H. C. Hotchkiss, G. J. Vermeij, K. R. Moonoosawmy, M. Huntley, B. Evans, B. Golding, and R. Mooi for comments on earlier versions of the manuscript. The Ontario Ministry of Training, Colleges, and Services and Natural Sciences and Engineering Research Council of Canada (Discovery Grant 261590) for financial support. MAC also thanks N. Abou Chakra, Arne Traulsen, and the Max Planck Society for their help and support.

## Authors Contributions Statement

MAC devised and developed the model, wrote the program, analysed the results and created the cdf module (available upon request). MAC measured specimens and analysed the results. ML conducted the analytical solution. MAC, ML, and JS contributed to the conceptualization of the catenary model. All authors reviewed the manuscript.

---

**Figure S1.**
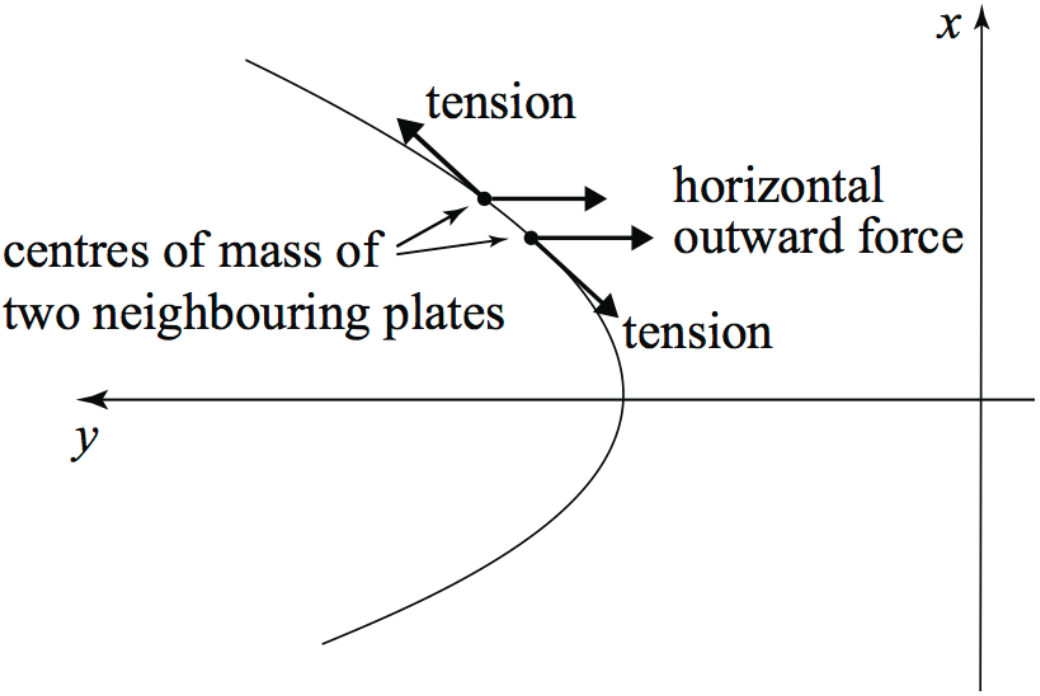
A diagram showing forces acting on a test: the tension force pulling apart neighbouring plates, and the outward-facing horizontal force, which approximately balances all other forces present.

**Figure S2.**
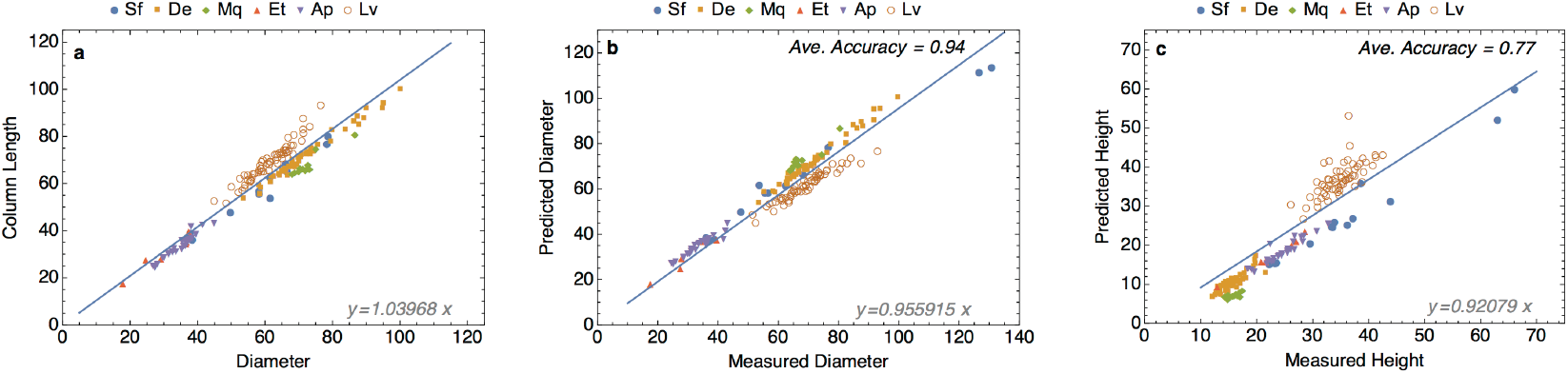
Data for specimens representing echinoid taxa: *S. franciscanus* (Sf), *D. excentricus* (De), *M. quinquiesperforata* (Mq), *E. thouarsii* (Et) *A. punctulata* (Ap), and *L. variegatus* (Lv). a) Measured column length vd measured diameter. b) Measured versus predicted diameter values. c) Measured versus predicted height values. Measurements are millimetres.

